# spatialMET: an open and scalable framework for spatial metabolomics analysis

**DOI:** 10.64898/2026.08.27.747606

**Authors:** Yonatan Ayalew Mekonnen, Oscar E. Ospina, Vanessa Rubio, Eric Welsh, Reaz Uddin, Hayley D. Ackerman, Alex Soupir, James E. Cox, Brooke L. Fridley, Elsa R. Flores, John Koomen, Paul A. Stewart

**Author notes:** These Authors contribute equally and share the first authorship (Yonatan Ayalew Mekonnen, Oscar E. Ospina).

## Abstract

Mass spectrometry imaging (MSI) enables spatially resolved metabolomics in intact tissue sections, but analysis remains challenging at scale. Existing MSI workflows often require users to combine multiple software tools, while others rely on proprietary vendor software that limits interoperability and reproducibility. To address these challenges, we developed spatialMET, an open-source framework that provides an end-to-end workflow for MSI analysis. spatialMET provides a unified platform for preprocessing, spatial domain detection, and visualization. Downstream analyses include differential abundance testing, spatial autocorrelation and gradient analysis, dimensionality reduction, and correlation network analysis. Spatial domain detection uses hcdist, a C-based hierarchical clustering implementation that substantially reduces runtime and memory use relative to existing R-based approaches. spatialMET can be run through an interactive R Shiny application or as a standalone command-line workflow for larger datasets or high-performance computing environments. Applied to mouse small cell lung cancer MALDI-MSI data containing 284,673 pixels, spatialMET identified tumor-associated, stromal, and adjacent lung spatial domains that aligned with matched histology. Differential abundance analysis identified 117 m/z features that differed between tumor and stromal regions, while spatial autocorrelation analyses revealed spatially structured abundance patterns. Applying spatialMET to mouse lung adenocarcinoma data from an entire lung lobe containing 338,477 pixels further demonstrated scalability and captured spatial heterogeneity across tumor and surrounding lung tissue. In summary, spatialMET provides a scalable, open-source framework for end-to-end spatial metabolomics analysis, and it is distributed as a Docker container for reproducible deployment. Source code and installation instructions are available at https://github.com/biodatalab/spatialMET.

## Introduction

Mass spectrometry-based metabolomics has identified biomarkers and metabolic alterations across many cancer types^1,2^. However, conventional metabolomics loses the spatial information needed to understand how metabolic activity varies across tissue regions. In cancer, metabolic programs vary across tumors and within individual tissues because of differences in tumor cell states, the local microenvironment, and interactions with surrounding stromal and immune cells^2^. Consequently, preserving spatial information is critical for understanding how metabolism is organized within tissues. Recent advances in spatial molecular profiling have shown that incorporating spatial information enables the identification of region-specific biochemical programs, improves interpretation of tissue heterogeneity, and links molecular distributions to histological and functional organization^3,4^. In parallel, developments in mass spectrometry imaging (MSI) now allow mapping and quantification of hundreds to thousands of metabolite-associated m/z features within tissue sections at high spatial resolution^4^.

Unlike conventional metabolomics, MSI preserves the spatial location of molecular measurements within tissue. This creates a need for analytical tools that can relate metabolite abundance to tissue architecture, spatial organization, and regional heterogeneity. Several tools have been developed for MSI analysis, including EXIMS^5^, Cardinal^6^, and SpaMTP^7^, which support data import, preprocessing, peak selection, visualization, and clustering. Although these tools support important steps in MSI analysis, many workflows still require users to combine multiple software packages to complete an analysis. This fragmentation can make downstream spatial analyses difficult to implement, particularly when additional programming or statistical expertise is required. Spatial statistics are widely used in spatial transcriptomics and other imaging modalities, but they remain less commonly integrated into MSI workflows. In addition, some analysis suites are restricted to proprietary vendor software, limiting interoperability and reproducibility. To address these limitations, we developed spatialMET, an open-source framework for spatial metabolomics analysis (Figure 1). Source code and installation instructions are available on GitHub (https://github.com/biodatalab/spatialMET).

**Figure 1.**
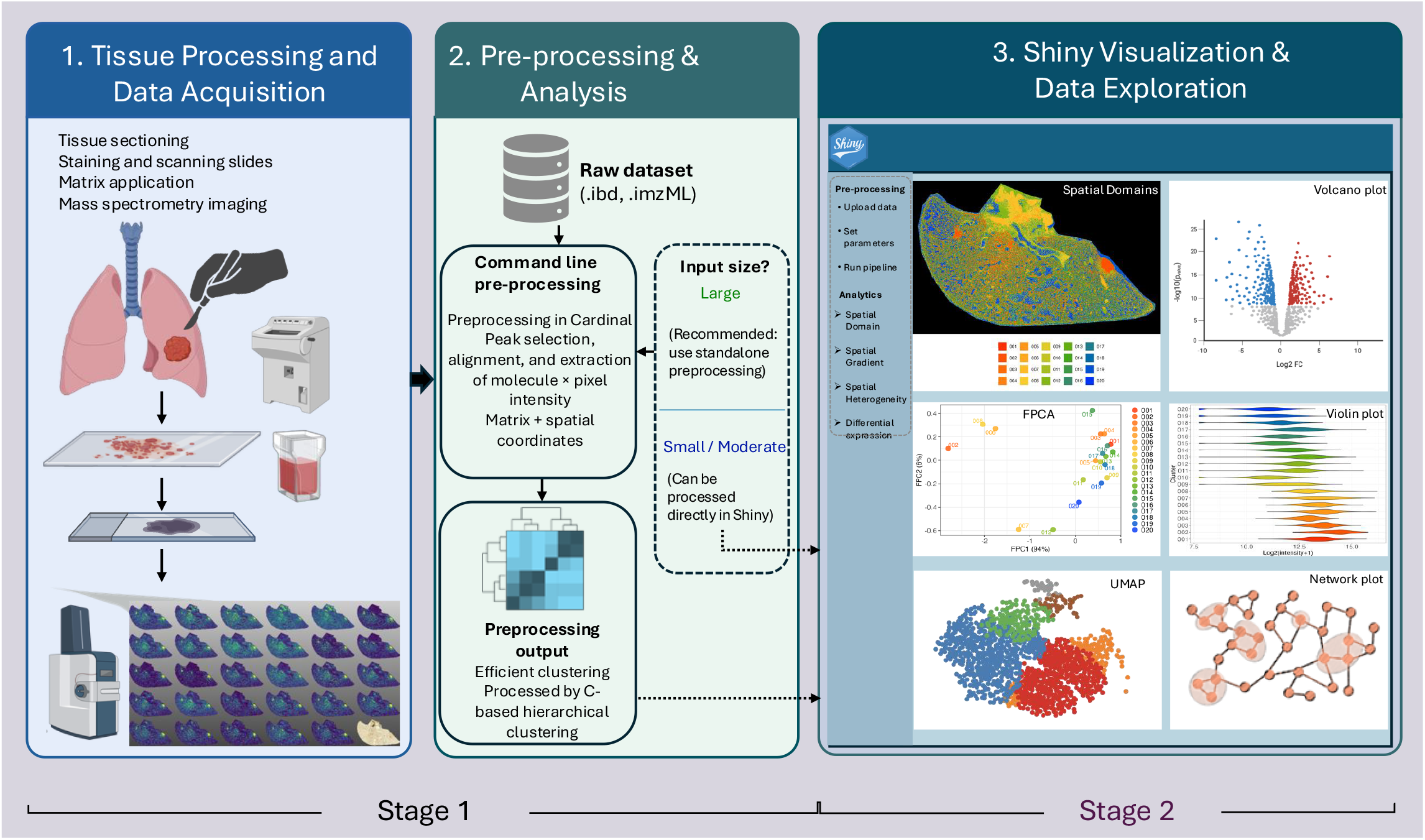
Overview of the spatialMET workflow. Raw mass spectrometry imaging (MSI) data (imzML and ibd files) are uploaded directly into the spatialMET Shiny application. For datasets of reasonable size, the application performs the complete preprocessing workflow, including peak detection, alignment, generation of molecule by pixel intensity matrices with spatial coordinates, filtering, and hcdist based hierarchical clustering to identify spatial domains. For larger datasets, the same preprocessing workflow can be performed externally using the standalone preprocessing pipeline before importing the processed data into the application. The resulting processed data are then used within the Shiny interface for interactive visualization, exploratory analysis, and downstream spatial analyses.

spatialMET combines computationally efficient preprocessing and spatial domain detection with an interactive suite of downstream analysis tools. Users can load raw MSI files (.imzML/.ibd) directly into the R Shiny application for peak filtering, data transformation, and hierarchical clustering of pixel-level m/z feature intensities. Processed MSI data and spatial domain assignments can then be analyzed in the same interface using a suite of tools for visualization, differential abundance testing, spatial autocorrelation analysis using Moran’s I^8^ and Geary’s C^9^, spatial gradient analysis, dimensionality reduction, and network analysis. For larger datasets or high-performance computing environments, the preprocessing and clustering steps can also be run from the command line, with outputs imported into the Shiny application for downstream analysis.

## Results – development and implementation of spatialMET

### Data preprocessing and input formats

spatialMET uses the Cardinal package^6^ for MSI data import, organization, and preprocessing. The framework accepts two input formats: 1) imzML and ibd files, the standard formats for MSI^10^, and 2) tab-delimited matrices containing peak intensities for each pixel along with corresponding spatial coordinates. These preprocessing steps can be run either from the command line for high-performance computing environments or within the R Shiny application for interactive analysis.

### Novel hierarchical clustering implementation

The resulting tab-delimited output files are processed using hcdist, a novel C-based hierarchical clustering implementation for tissue domain detection. hcdist generates a dendrogram that is cut using a user-specified number of clusters to assign spatial domain labels to individual pixels. Pixels are then plotted in their original X-Y coordinates with domain-specific colors, allowing users to visualize spatially coherent metabolic regions and m/z feature intensities across tissue. hcdist improves computational performance through a multithreaded C implementation that parallelizes distance matrix calculations and hierarchical clustering node updates, together with a novel seriation heuristic for optimizing node-flipping.

### Shiny application and interactive visualization

Hierarchical clustering with hcdist can be executed either through the R Shiny application or through the standalone command-line workflow for larger datasets and computing cluster environments. Following clustering with hcdist, pixel-level domain assignments and peak intensity matrices are imported into the Shiny application for visualization and downstream analysis. Users can also upload a matched hematoxylin and eosin (H&E)-stained histology image in PNG, JPEG, or TIFF format to compare molecular patterns with tissue morphology. The application includes modules for spatial domain visualization, m/z feature intensity mapping, differential abundance analysis, spatial statistics, PCA, UMAP embedding, and spatial gradient analysis. Additional visualization tools generate volcano plots, violin plots, and scatter plots to characterize m/z feature abundance across tissue regions.

### Spatial gradient analysis

Building on the spatial domain assignments, the spatial gradients module adapts the STgradient^11^ algorithm, originally developed for spatial transcriptomics, to identify m/z features exhibiting spatial abundance gradients relative to tissue domains, including features enriched at domain interfaces or boundary regions. The module uses cluster assignments, spatial coordinates, and abundance values to identify features that increase or decrease with distance from a user-selected reference domain. For each m/z feature, linear and rank-based (Spearman) associations between m/z feature intensity and distance from the reference domain are tested, yielding slopes, p-values, and FDR-adjusted values for m/z features with spatial gradients across the tissue.

### Differential abundance and spatial statistics

The differential abundance module supports multiple approaches for testing abundance differences between spatial domains, including Wilcoxon’s rank-sum test, Student’s t-test, and spatial limma^12,13^. In addition, Hellinger distance^14^ is calculated to quantify compositional dissimilarity between spatial regions, allowing comparison of overall m/z feature profiles across tissue domains. The spatial statistics module calculates spatial autocorrelation in m/z feature abundance patterns using Moran’s I^8^ (range: −1 to +1; positive values indicate spatial clustering, values near 0 indicate random spatial distribution, and negative values indicate dispersion) and Geary’s C^9^ (range: 0 to 2; values <1 indicate clustering, values near 1 indicate random spatial distribution, and values >1 indicate dispersion). Spatial weights are constructed using a k-nearest neighbor approach, and Moran’s I and Geary’s C are used to quantify spatial clustering of m/z feature intensities. These statistics help identify m/z features with spatially coordinated abundance patterns, including localized hotspots or dispersed distributions, that may reflect tissue architecture or microenvironmental organization.

### Dimensionality reduction and m/z feature abundance distributions

The Shiny application supports exploratory analysis of m/z feature variability using PCA, UMAP, and distribution plots. PCA is performed using the default R implementation, and UMAP is implemented through the uwot package. PCA and UMAP embeddings can be visualized with spatial domain annotations to examine variation in pixel-level m/z feature profiles across spatial domains. UMAP parameters, including the number of neighbors and minimum distance, can be specified by the user. m/z feature intensity distributions can also be compared across spatial domains using violin plots, boxplots, or both, with optional log2 transformation and colors matched to the spatial domain assignments.

The Shiny application also supports functional data analysis (FDA)^15^ to summarize spatial patterns that vary continuously across tissue space. In this approach, m/z feature profiles are represented as functions rather than as isolated pixel-level measurements or domain-level summaries. Functional principal component analysis (FPCA) is then used to identify major patterns of variation in these spatial profiles, allowing m/z features or tissue regions to be compared based on shared spatial organization.

### Domain-specific abundance and co-localization analysis

To summarize domain-level abundance patterns, the top N most variable m/z features are selected across all pixels based on variance. For each spatial domain, the median intensity of each m/z feature is compared with its median intensity across all remaining pixels, and domain-specific abundance is quantified as log2 fold change (log2FC). m/z features are considered elevated within a domain using a user-defined fold-change threshold, such as log2FC > 1, corresponding to at least 2-fold higher median abundance. The number of elevated m/z features per domain is summarized in a table and visualized as a bar plot.

To evaluate co-localization patterns, Pearson or Spearman correlation coefficients are computed between m/z feature intensities within a selected domain or across the full tissue section^16^. Network edges are established between m/z feature pairs whose correlation exceeds a user-defined threshold, and the resulting network is visualized interactively to highlight co-varying spatial abundance patterns. In this representation, nodes represent individual m/z features, while edges denote co-variation in spatial abundance. Positive correlations suggest co-localization, indicating m/z features that tend to appear together in the same tissue regions. These patterns may reflect shared spatial localization or related biochemical processes, although correlation alone does not establish a direct biochemical relationship. In contrast, negative correlations indicate mutually exclusive spatial distribution patterns.

## Results - Benchmarking

### Spatial domain detection benchmarking

To assess spatial domain detection, we compared hcdist with spatial shrunken centroids (SSC), a widely used method implemented in Cardinal v3.6.5. To evaluate agreement across clustering resolutions, hcdist and SSC were run with K = 10, 15, and 20 domains. hcdist was also compared with SSC across sparsity parameters s = 0, 3, 6, 9, 12, and 15. For all SSC runs, the neighborhood radius parameter r was set to 0 so that neighboring pixel information was not used during clustering. Agreement between hcdist and SSC domain assignments was evaluated using the Adjusted Rand Index (ARI; flexclust v1.4.2), which measures similarity between clustering solutions while accounting for agreement expected by chance^17^. ARI values ranged from 0.250 to 0.314, with the highest agreement observed for K = 10 (ARI: 0.277–0.314), followed by K = 15 (ARI: 0.273–0.299) and K = 20 (ARI: 0.250–0.299). The lower ARI values at higher K may reflect increased sensitivity to background noise and fine-scale domain boundaries as increasing K produces finer spatial partitions (Figure S1).

### hcdist performance benchmark

We benchmarked hcdist against fastcluster and parallelDist for hierarchical clustering using a 320 MB matrix containing 51,196 pixels and 2,107 m/z features. On a 64-core system, hcdist completed distance matrix computation and hierarchical tree construction in 3 minutes and 18 seconds while using 10.13 GB of memory (Figure 3G). These results demonstrate substantially improved runtime and memory use relative to existing R-based hierarchical clustering implementations.

### Resource utilization scaling across feature levels

To evaluate scalability with increasing feature counts, we measured resource utilization across six feature-retention levels using a 5 GB small cell lung cancer (SCLC) dataset: 100%, 75%, 50%, 25%, 10%, and 5% of m/z features. For each configuration, we assessed working RAM, private memory, and CPU utilization. Benchmarking was performed on a system equipped with a 13th Gen Intel® Core™ i5-13500H processor (12 physical cores, 16 logical processors; base frequency 2.60 GHz) and 8 GB RAM. Processing the complete dataset at the 100% feature level required 55 seconds, showing that the full feature set could be processed on a laptop-class system with limited memory. The 100% feature level had the highest peak working RAM (1.5 GB) and peak private memory (4.3 GB), reflecting the increased demand of processing the complete feature set, whereas average working RAM and average private memory remained relatively stable across feature-retention levels. These results indicate that higher feature counts primarily affected peak memory demand rather than sustained memory use. CPU utilization was also consistent across feature-retention levels, with similar peak and average use across analyses (Figure S2).

### Comparison with existing spatial metabolomics platforms

We compared spatialMET with existing spatial metabolomics platforms, including Cardinal^6^, Multi-MSIprocessor^18^, SMAnalyst^19^, and SpaMTP^7^. As summarized in Table S1, these tools support key steps such as preprocessing, visualization, and clustering, but few platforms provide integrated support for spatial autocorrelation, gradient analysis, multi-sample domain detection, and downstream association testing with experimental or clinical variables. These analyses often require manual scripting or external software. spatialMET combines preprocessing, spatial domain detection, spatial autocorrelation analysis, gradient analysis, and differential abundance testing within a single graphical workflow. The accompanying R Shiny interface provides interactive visualization and downstream analysis with reduced need for custom scripting.

### Ion Mobility Mass Spectrometry Imaging

Intact lung lobes from small cell lung cancer (SCLC) and non-small cell lung cancer (NSCLC) mouse models were prepared for matrix assisted laser/desorption ionization-ion mobility-mass spectrometry imaging (MALDI-IM-MSI) using a previously published protocol^20^. Briefly, murine lungs were insufflated with Milestone Cryoembedding Compound (MCC; Milestone Medical, 6705MILE01) and immediately snap frozen in liquid nitrogen (LN2). Frozen lung tissues were cryosectioned at 5 μm thickness and mounted either on standard microscopy slides for H&E staining and immunofluorescence or on indium-tin oxide (ITO) coated slides for MALDI-IM-MSI. ITO slides were sprayed with N-(1-naphthyl) ethylenediamine dihydrochloride (NEDC) (Tokyo Chemical Industry, N0869) at 7 mg/mL in aqueous 50% MeOH using an HTX M5 Sprayer. Data were acquired on a Bruker timsTOF fleX MALDI ion mobility mass spectrometer rastering at 20 μm raster step size across the entire lung tissue sections. Imaging data were exported using SCiLS Lab (v. 12.01 Core, Bruker) to the open imzML format.

## Results – case studies in mouse lung cancer

We applied spatialMET to mouse small cell lung cancer (SCLC) tissue sections to evaluate its ability to resolve spatial metabolic organization from MSI data (Figure 2). The matched H&E-stained section showed the underlying tissue architecture (Figure 2A), and spatialMET domain assignments captured corresponding regional organization in the MSI data (Figure 2B). Higher-magnification views showed distinct spatial patterns within stromal and adjacent lung regions (Figure 2C, D) and multiple smaller domains within tumor-associated regions (Figure 2E–G). The presence of multiple tumor-associated domains is consistent with intratumoral heterogeneity in m/z feature abundance. Together, these findings show that spatialMET can identify tissue-associated spatial domains and characterize regional metabolic heterogeneity directly from MSI data (Figure S3).

**Figure 2.**
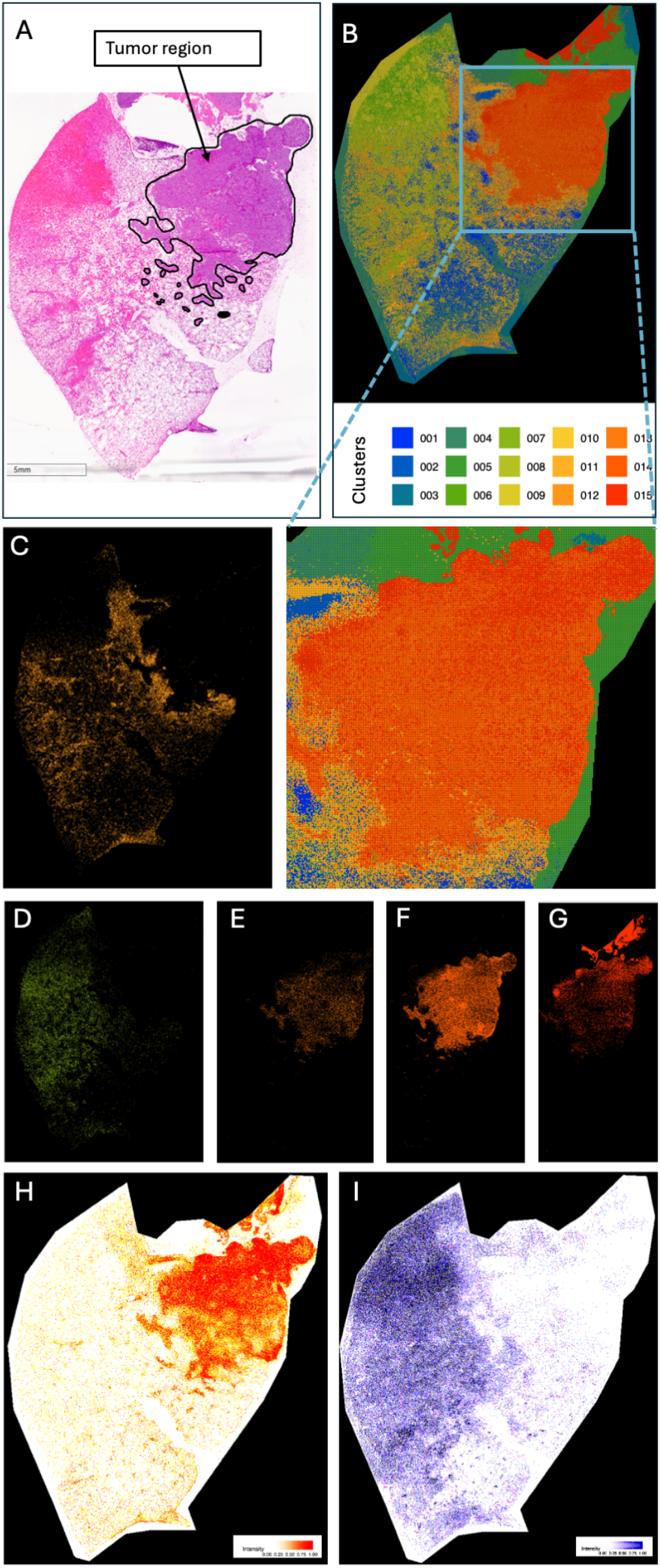
spatialMET identifies tumor-associated spatial domains in small cell lung cancer (SCLC). (A) Hematoxylin and eosin (H&E)-stained tissue section showing histologically defined tumor regions. (B) spatialMET-inferred spatial domains reveal coherent tumor-associated clusters, with a zoomed inset highlighting a representative domain with distinct spatial boundaries and intradomain heterogeneity. (C,D) Spatial distributions of stromal- and adjacent tissue-associated domains. (E-G) Spatial distributions of tumor-associated domains identified by spatialMET. (H) Spatial distribution of the tumor-associated m/z feature 794.5669. (I) Spatial distribution of the stromal-associated m/z feature 471.2578.

We identified 50 m/z features exhibiting strong spatial autocorrelation (Moran’s I ≥ 0.6 and Geary’s C ≤ 0.3). Among them, m/z 794.5669 showed localized abundance in tumor-associated regions (Moran’s I = 0.78, Geary’s C = 0.22; Figure 2H). In contrast, m/z 471.2578 showed a broader spatial distribution (Moran’s I = 0.62, Geary’s C = 0.38; Figure 2I). Differential abundance analysis using the Wilcoxon rank-sum test identified 117 m/z features that differed significantly between tumor and stromal regions (adjusted p < 0.05 and absolute log2 fold change ≥ 1; Figure 3A), showing that tumor and stromal domains differed in m/z feature abundance.

UMAP analysis showed structured variation in pixel-level m/z feature profiles across the tissue section (Figure 3B). FPCA provided a complementary summary of spatial m/z feature variation, separating m/z features according to major patterns of abundance across tissue domains (Figure 3F). Overall domain composition and heterogeneity are summarized in Figure 3C. Across adjacent, stromal, and tumor regions, the relative abundance of variable domains shifted, showing regional differences in m/z feature abundance across the tissue microenvironment (Figure 3D). The top 30 most variable m/z features across the tissue section were used for correlation network analysis, which identified 19 highly correlated features (|r| ≥ 0.6) with similar abundance patterns within tumor-associated regions (Figure 3E).

**Figure 3.**
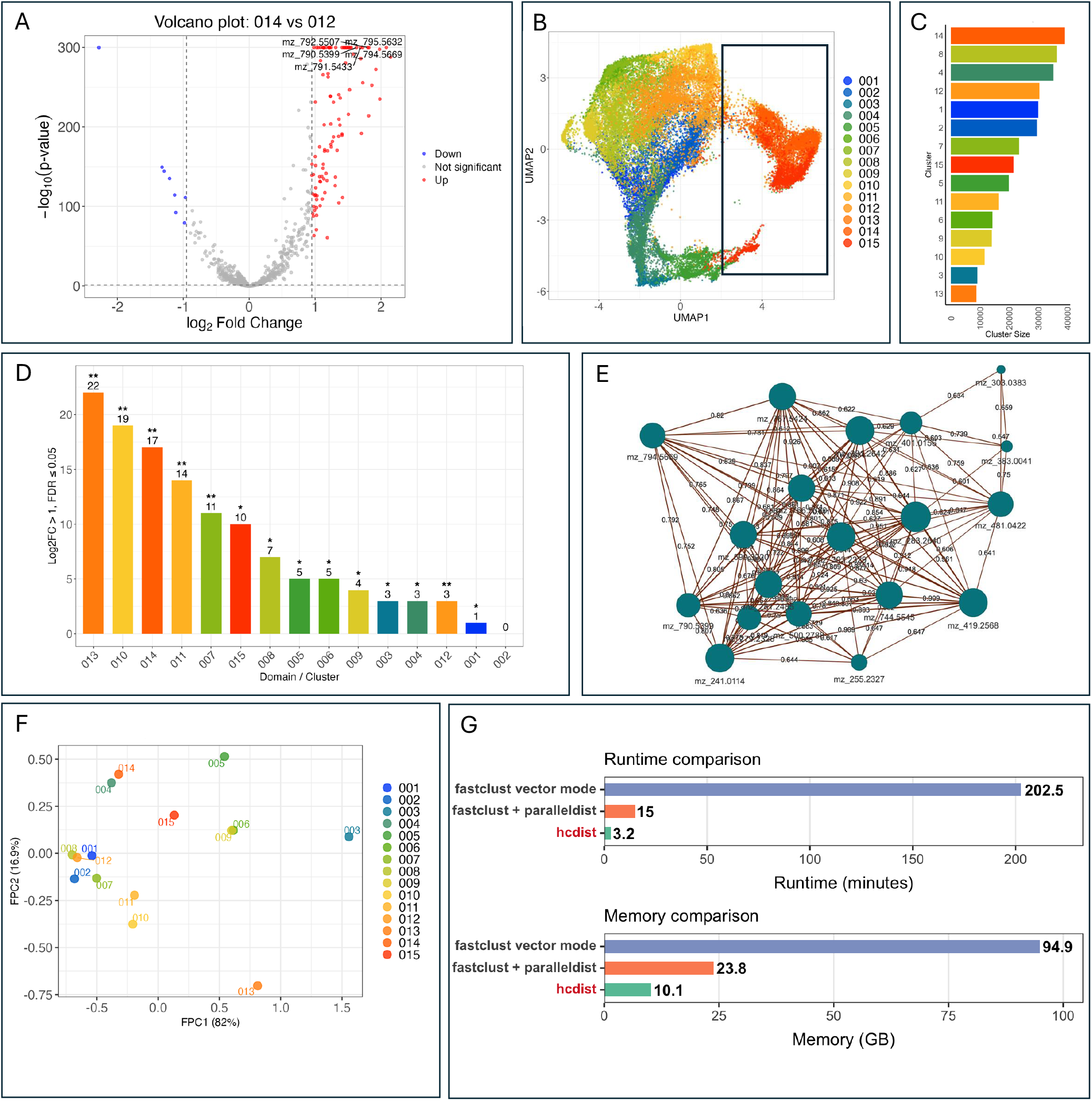
Integrated spatial analyses reveal metabolic organization, spatial dependence, and tumor-domain structure across samples, together with computational benchmarking. (A) Volcano plot of differentially abundant m/z features across spatial domains, highlighting significantly increased and decreased features. (B) UMAP projection showing metabolically distinct clusters across spatial domains, with representative regions highlighted. (C) Domain composition analysis illustrating spatial heterogeneity across samples. (D) Bar plot summarizing the number of m/z features with domain-specific abundance patterns, defined by absolute log_2_ fold change ≥ 1 and false discovery rate (FDR) ≤ 0.05. (E) Correlation network constructed from the 30 most variable m/z features across tumor regions. Nodes represent m/z features, with node size proportional to network degree; edges indicate pairwise correlations above the selected threshold (|r| ≥ 0.6). (F) Functional principal component analysis of spatial profiles, revealing structured spatial variation across samples. (G) Runtime and memory benchmarking of hierarchical clustering methods on a 320 MB dataset.

We next applied spatialMET to a mouse lung adenocarcinoma specimen to evaluate performance in a second lung cancer histology. spatialMET identified multiple spatial domains across the tissue section, including tumor-associated and microenvironmental regions. Representative spatial footprints showed organized m/z feature abundance patterns in a histology distinct from the SCLC specimen (Figure S4, Figure S5).

## Conclusions

spatialMET combines computationally efficient spatial domain detection with an interactive R Shiny application for spatial metabolomics analysis. Core preprocessing and clustering steps can be executed either directly within the Shiny application or, for large-scale datasets, through a standalone command-line workflow suited to high-performance computing environments. The workflow supports data preprocessing, spatial domain detection, m/z feature intensity visualization, differential abundance testing, spatial autocorrelation analysis, and spatial gradient detection. Applied to lung cancer MALDI-MSI datasets, spatialMET identified spatial domains corresponding to tumor-associated, stromal, and adjacent lung regions, with patterns that aligned with matched histologic annotations. Benchmarking showed substantially reduced runtime and memory use compared with existing approaches. By integrating preprocessing, spatial clustering, statistical analysis, and interactive visualization, spatialMET brings key MSI analysis steps into a single workflow. Overall, spatialMET provides an open-source tool for interpreting spatially organized m/z feature abundance patterns and extending spatial metabolomics analyses to multimodal tissue studies.

## Author contributions

YAM, OEO, JK, and PAS conceptualized and designed the study. YAM led the development of the spatialMET software, integrated the C-based hcdist clustering algorithm with the Shiny application, implemented the preprocessing pipeline, developed the interactive visualization modules, and built the Docker container for reproducible deployment. YAM wrote the first draft of the manuscript and created all figures. OEO co-developed the software and contributed to the Shiny app design. EW provided code for the data preprocessing steps (hcdist). VR contributed to data generation. YAM and OEO performed the data analysis, with support from RU, AS, and HA. JEC, BLF, and ERF contributed to funding acquisition, investigation, and resources. PAS, JK, and ERF co-supervised the study. All authors reviewed and approved the final manuscript.

## Acknowledgements

This research was funded by the Biostatistics and Bioinformatics Shared Resource, and the Proteomics and Metabolomics Shared Resource at the H. Lee Moffitt Cancer Center & Research Institute through the support of the Cancer Center Support Grant (P30 CA076292), the Cancer Research Institute Technology Impact Award, P01 CA250984 (“Identifying Metabolic Vulnerabilities in Lung Cancer”), and the Fulbright Scholar Program. The research reported in this publication was supported by Huntsman Cancer Foundation and the National Cancer Institute of the National Institutes of Health under Award Number P30 CA042014. This publication has been supported by funding from the National Institutes of Health (NIH). Any opinions, findings, and conclusions or recommendations expressed in this material are those of the author(s) and do not necessarily reflect the views of the NIH.

