## Supplemental Figure 1 for "spatialMET: an open and scalable framework for spatial metabolomics analysis"

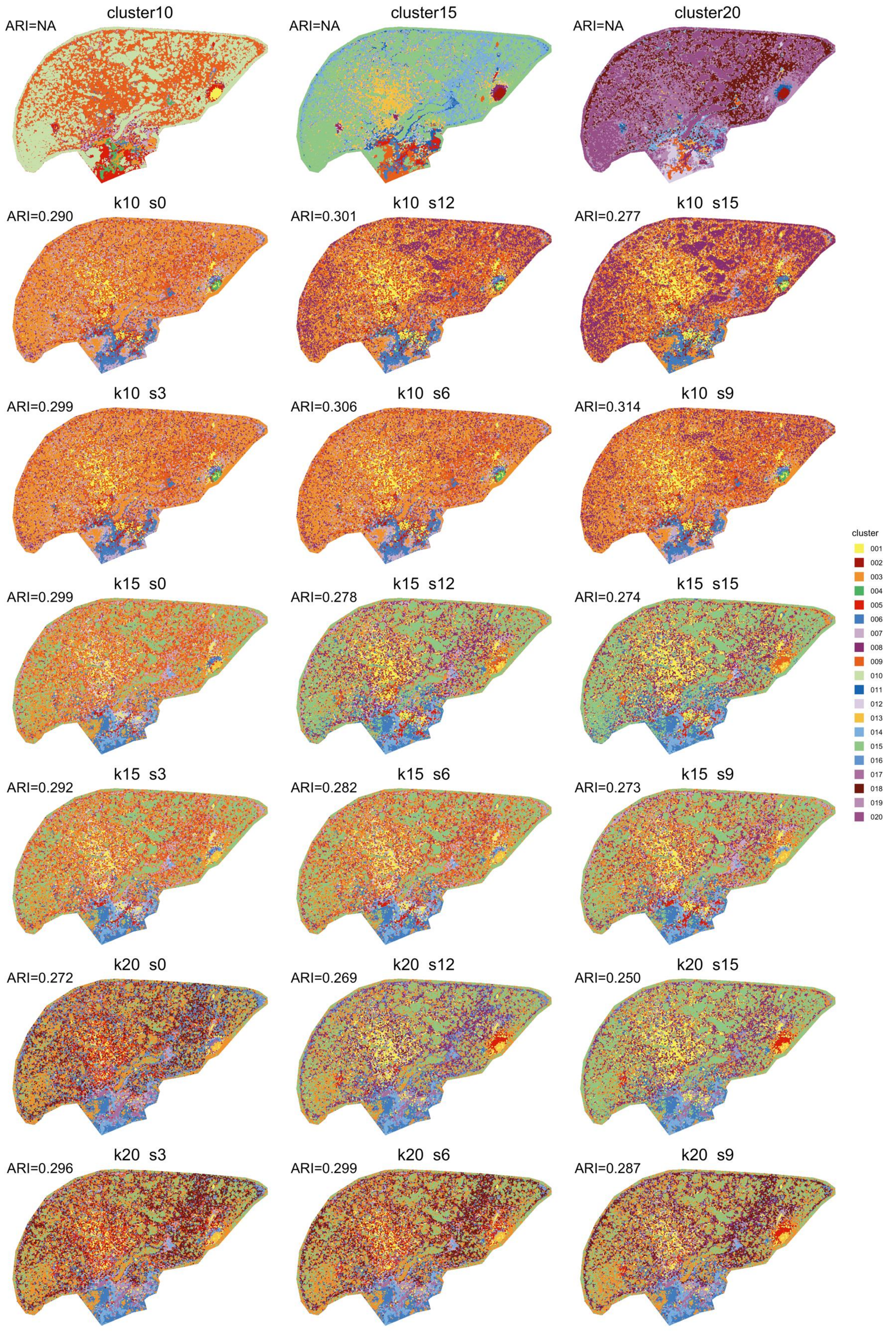


**Figure S1. Comparison of pixel-level spatial domain assignments from hcdist and spatial shrunken centroids (SSC).** Pixel-level spatial domain assignments are shown for hcdist and SSC across K values of 10, 15, and 20 and SSC sparsity parameters of s = 0, 3, 6, 9, 12, and 15. The adjusted Rand index (ARI) was calculated to measure agreement in spatial domain assignments between the two methods.

.
