## Supplemental Figure 2 for "spatialMET: an open and scalable framework for spatial metabolomics analysis"

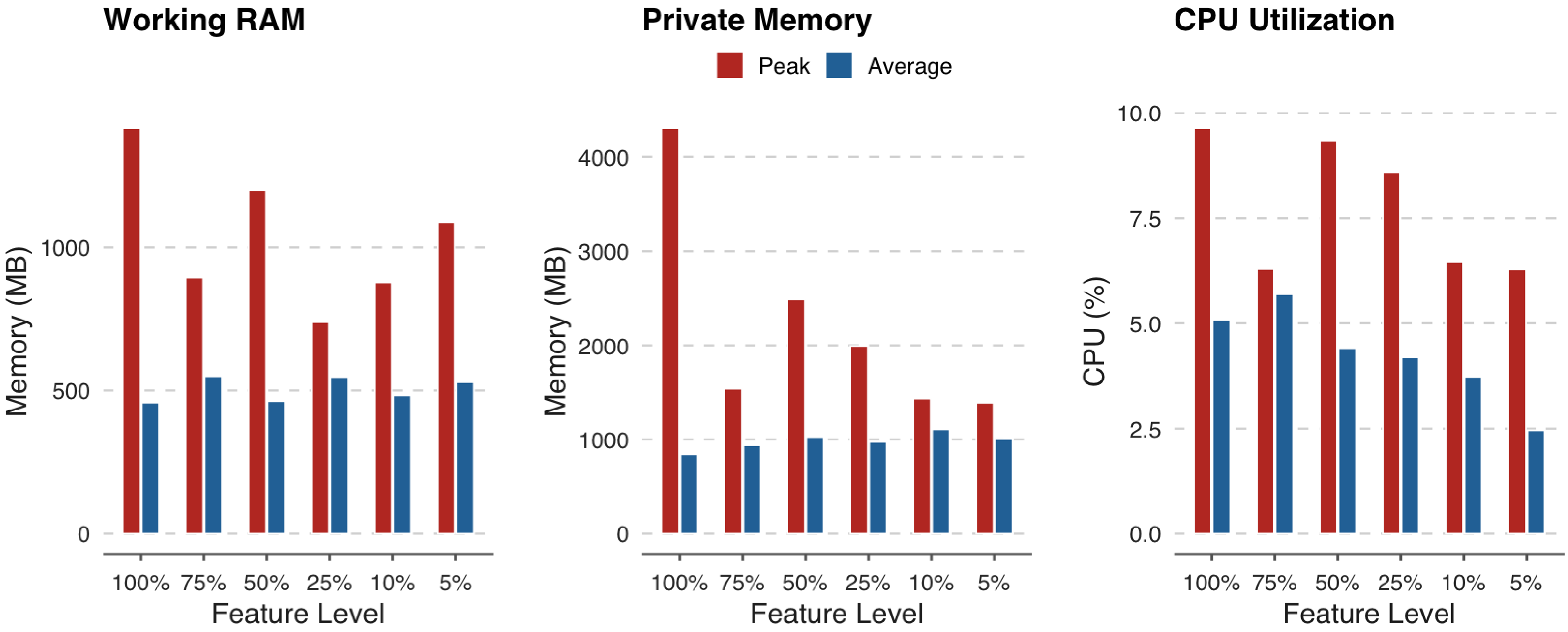


**Figure S2. System resource utilization across feature-retention levels.** Peak and average working RAM, private memory, and CPU utilization were recorded for six feature-retention levels (100%, 75%, 50%, 25%, 10%, and 5%) using a 5 GB dataset. Peak values are shown in red and average values in blue across all panels.
