## Supplemental Figure 3 for "spatialMET: an open and scalable framework for spatial metabolomics analysis"

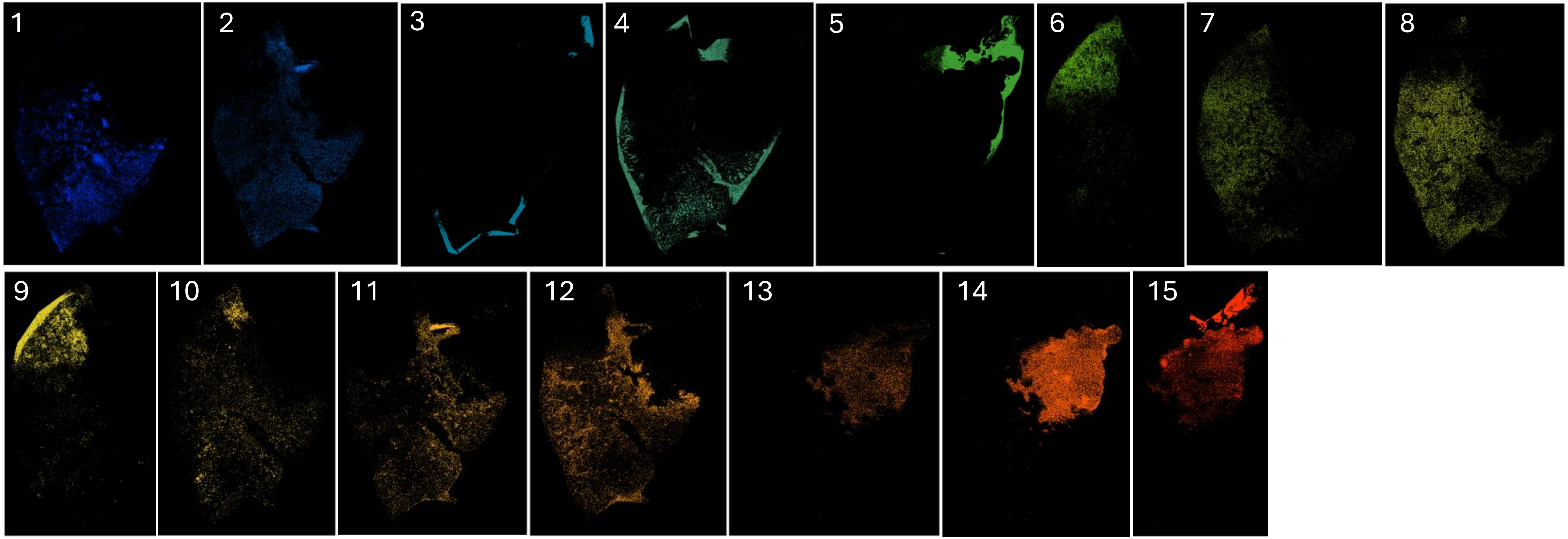


**Figure S3. Spatial footprints of representative clusters identified across the full SCLC tissue section.** Each image shows a single cluster, with all other tissue regions removed, to depict only the pixels assigned to that cluster. This view highlights the spatial distribution of individual clusters and enables visualization of microenvironmental regions within the SCLC tissue, including tumor mass, invasive margin, stromal compartments, and necrotic regions.
