## Supplemental Figure 4 for "spatialMET: an open and scalable framework for spatial metabolomics analysis"

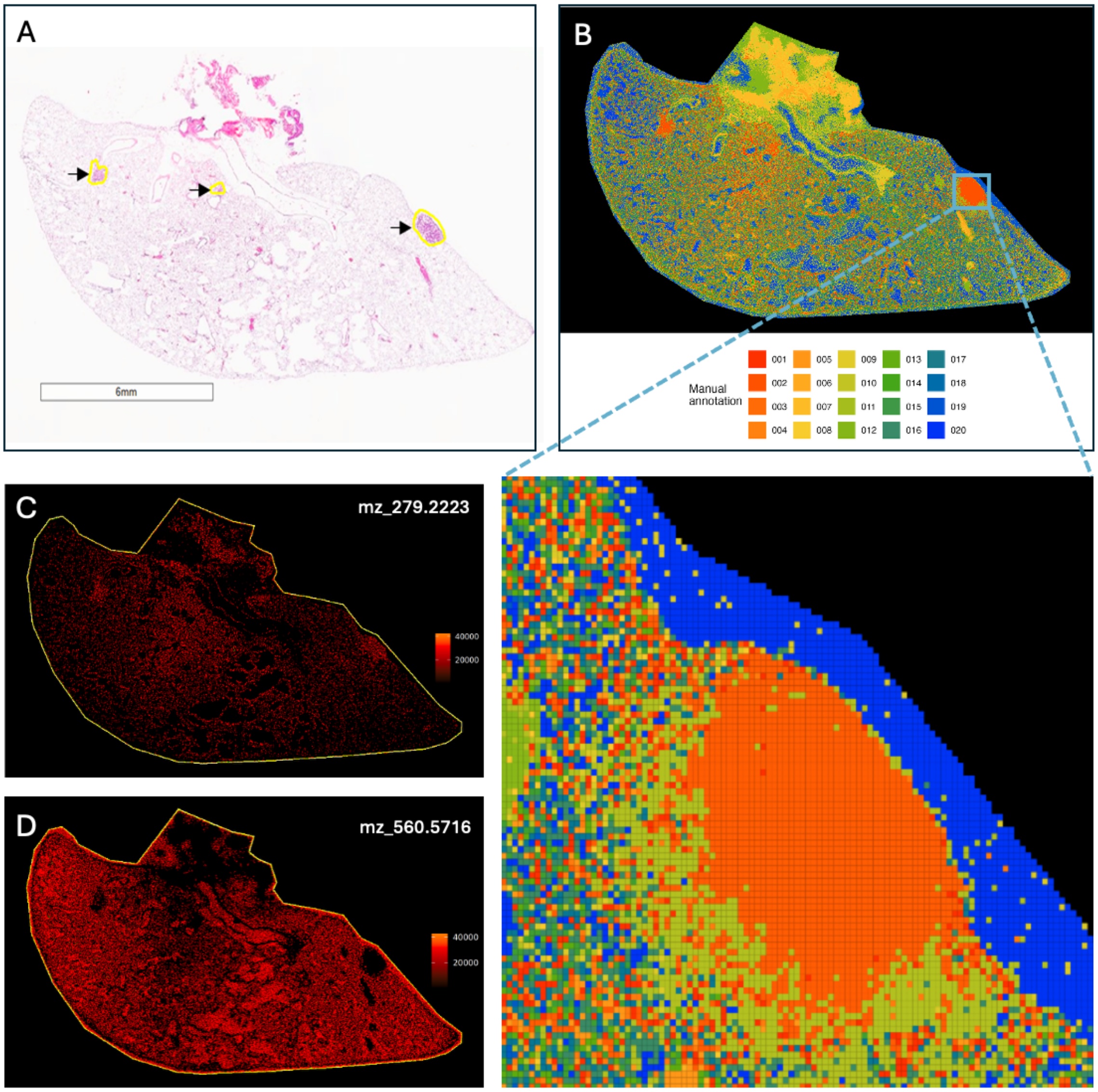


**Figure S4. spatialMET identifies tumor-associated spatial domains in non-small cell lung cancer (NSCLC).** (A) Hematoxylin and eosin (H&E)-stained tissue section showing histologically defined tumor regions. (B) spatialMET-inferred spatial domains reveal coherent tumor-associated clusters, with a magnified inset highlighting representative domains with distinct spatial boundaries and intradomain heterogeneity. (C) Spatial distribution of m/z feature 279.2223, demonstrating gradual molecular transitions across spatial domains. (D) Spatial distribution of m/z feature 560.5716, illustrating intradomain molecular uniformity within spatialMET-identified regions.
