## Supplemental Figure 5 for "spatialMET: an open and scalable framework for spatial metabolomics analysis"

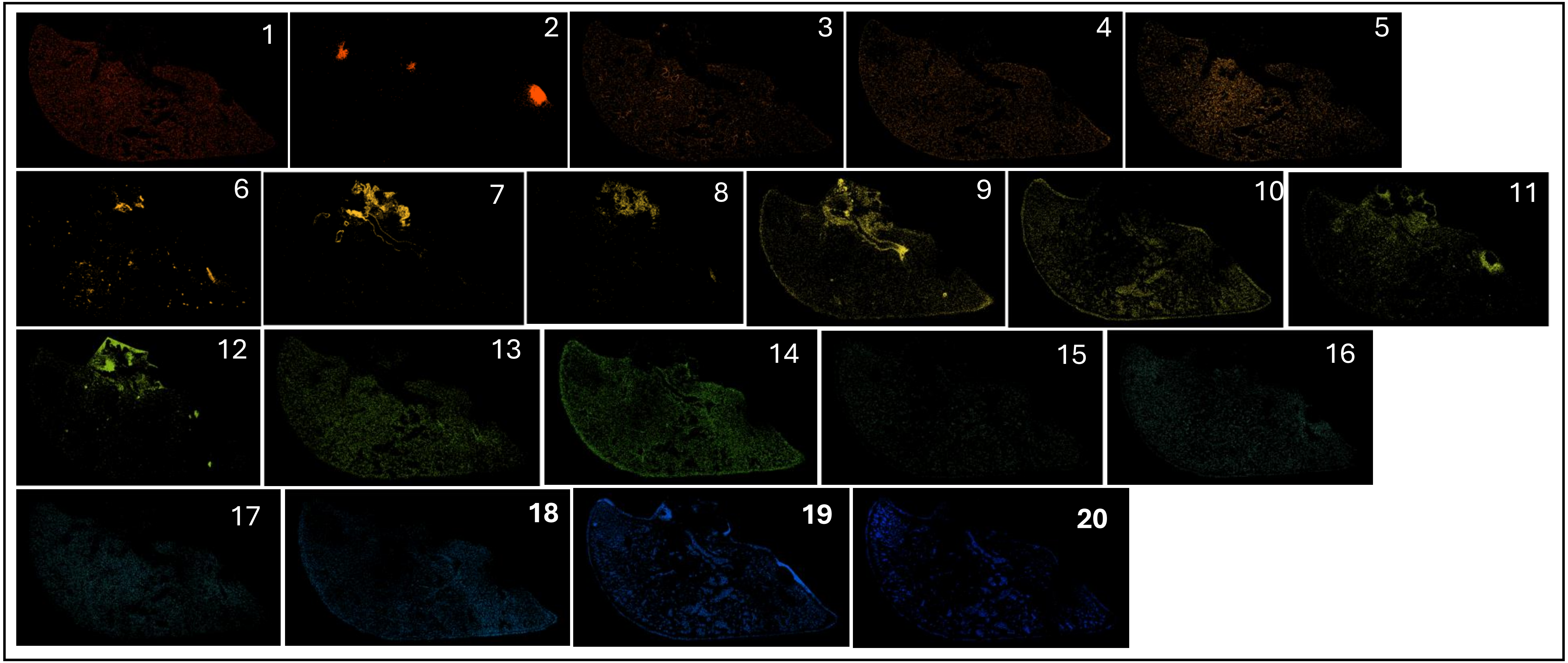


**Figure S5**. Spatial footprints of representative clusters identified throughout the full NSCLC tissue section. Each image illustrates a single cluster and removes all other areas of tissue, using a uniform cluster specific color scheme, to depict only the pixels assigned to that cluster. Focusing on the spatial distribution of each identified cluster allows visualization of various microenvironmental niches within the SCLC tissue, such as the tumor mass, invasive margin, stromal compartments, and necrosis.
