## Supplemental Table 1 for "spatialMET: an open and scalable framework for spatial metabolomics analysis"

**Table S1.** Comprehensive Comparison of Spatial Metabolomics Software: Benchmarking and Functional Feature Assessment

| Category / Metric | spatialMET | Cardinal | Galaxy-MSI | SpaMTP | SpectralAnalysis | Multi-MSIprocessor | SMAnalyst | SMEW |
|---|---|---|---|---|---|---|---|---|
| **Data Ingestion Standard** | Native Raw (.imzML, .ibd) & Processed | Raw & Processed (.imzML, .analyze) | Raw & Processed (.raw, .mzML) | imzML, Cardinal Objects | imzML, .raw, .mzML | Raw Vendor Files | Processed Matrix only | Processed Tabular CSV only (Requires external scripts) |
| **End-to-End Workflow Integration** | Unified Native Pipeline (Raw to Stats) | Scripting required | Fragmented modules | Partial Cardinal dependency | GUI focused | Pre-processing only | Downstream only | Broken Workflow (Requires Stage-1 script pre-processing) |
| **Pre-processing & QC** | Feature/Pixel QC, Normalisation & Filtering | Baseline & Smoothing | Native Peak Filtering | Supported via Cardinal | Pixel-wise Smoothing | Basic Peak Filtering | Background QC | Native Spatial Denoising & Normalisation |
| **Dimensionality Reduction** | Integrated PCA & Spatial Coordinates | PCA, PLS | Not Natively Supported | PCA | PCA, NMF | Not Natively Supported | PCA, UMAP, t-SNE | PCA, PLS-DA, NMF, UMAP |
| **Clustering Capabilities** | High-Performance C-backend (hcdist) & K-means | Spatial Shrunken Centroids | Not Natively Supported | K-means, Hierarchical | Louvain, SLM | K-means | K-means (via Seurat) | K-means, Louvain, BayesSpace |
| **Differential Abundance** | Native Parametric/Non-Parametric Wilcoxon (Wilcoxon-test, spatial limma, and Hellinger distance) | Native Segmentation | Native t-tests | Native Wilcoxon (FDR) | Native Wilcoxon (FDR) | Not Available | Native Wilcoxon | Native Parametric & Non-Parametric Tests |
| **Tissue Annotation (ROI)** | Native Interactive Graphical Lasso Tool | Requires R Coding | Requires Manual Mapping | Lacks Native Selection | Lacks Native Selection | Lacks Native Selection | Lacks Native Selection | Native Interactive Selection & H&E Co-registration |
| **Spatial Gradients** | Native STgradient Module | Lacks Native Module | Lacks Native Module | Lacks Native Module | Lacks Native Module | Lacks Native Module | Lacks Native Module | Native Radial Distance Analysis |
| **Spatial Autocorrelation** | Native Moran's I & Geary's C | Requires Manual Coding | Lacks Spatial Module | Native SpaGene | Native Moran's I | Non-Spatial FELLA | Lacks Spatial Module | Native semla & MERINGUE |
| **Spatial Feature Selection** | Supported via Cardinal Integration | Native SSC Selection | Not Supported | Native SpaGene | Not Supported | Not Supported | Not Supported | Native Spatially Variable Peaks |
| **Biological Concordance / ST Validation** | External/Custom Integration | Custom R Integration | Not Supported | Native Seurat/ST Alignment | Not Supported | Not Supported | Not Supported | Native GENIE3 & MAGPIE ST Alignment |
| **Metabolite Feature Annotation** | Streamlined Local Reference Mapping (No API Setup) | Requires External Packages | HMDB/METLIN Integration | Internal HMDB/KEGG | Native m/z Matching | Integrated HMDB/METLIN | Not Natively Supported | Requires External Manual Script Setup via REST APIs |
| **FINAL SCORE** | **10 / 12** | **6 / 12** | **4 / 12** | **8 / 12** | **7 / 12** | **4 / 12** | **4 / 12** | **9/ 12** |

**Scoring Basis**: Each tool is evaluated across the 12 key operational categories. **Point Allocation Criteria (Binary Support Evaluation)**: **1 Point (Native/Integrated Support)**: Awarded when a feature is fully native, out-of-the-box, or integrated seamlessly within the tool's core execution environment (such as GUI-driven interactive lasso selection, native C-compiled backend algorithms, or unified raw-to-stats execution). This corresponds to the green-colored cells in the benchmark matrix. **0 Points (Non-Native/Code-Dependent/Absent)**: Assigned when a feature is either entirely missing, marked as "Lacks native modules", or when support requires external user intervention. This includes workflows described as "Requires manual R coding", "Requires manual coordinate mapping", "Requires external packages", "Requires Stage-1 script pre-processing", or "Requires External Manual Script Setup via REST APIs". This stringent penalty reflects the user-accessibility barrier for non-programmers.
